# Molecular Phylogenetic Analysis of *16S rRNA* Sequences Identified Two lineages of *H. pylori* Strains Detected from Different Regions in Sudan Suggestive of Differential Evolution

**DOI:** 10.1101/2019.12.25.888396

**Authors:** Abeer Babiker Idris, Hadeel Gassim Hassan, Maryam Atif Salaheldin Ali, Sulafa Mohamed Eltaher, Leena Babiker Idris, Hisham N. Altayb, Amin Mohamed Abass, Mustafa Mohammed Ahmed Ibrahim, El-Amin Mohamed Ibrahim, Mohamed A. Hassan

## Abstract

**Background:** *H. pylori* is ubiquitous among humans, and one of the best studied examples of an intimate association between bacteria and humans. Under several diverse socio-demographic factors in Sudan, a continuous increase in the prevalence rate of *H. pylori* infection has been noticed which represents a major public health challenge. In this study, we analyzed the molecular evolution of *H. pylori* Strains detected from different ethnic and regions of Sudan using *16S rRNA* gene and phylogenetic approach.

**Materials and methods:** A total of 75 gastric biopsies taken from patients who had been referred for endoscopy from different regions of Sudan. The DNA extraction was done by using the guanidine chloride method. Two sets of primers (universal and specific for H. pylori) were used to amplify the *16S ribosomal* gene. Sanger sequencing was performed; then Blast these sequences with those available in the NCBI nucleotide database. The evolutionary aspects were analyzed using a MEGA7 software.

**Result:** Molecular detection of *H. pylori* has shown that 28 (37.33%) of patients were positive for *H. pylori*. Bivariate analysis has found no significant differences exhibited across sociodemographic, endoscopy series and *H. pylori* infection. Nucleotide variations were found at five nucleotide positions (positions 219, 305, 578, 741 and 763-764) and one insertion mutation (750_InsC_751) was present in sixty-seven percent (7/12) of our strains. The phylogenetic tree diverged into two lineages.

**Conclusion:** The phylogenetic analysis of *16S rRNA* sequences identified two lineages of *H. pylori* strains detected from different regions in Sudan. Sex mutations were detected in regions of the *16S rRNA* not closely associated with either tetracycline or tRNA binding sites. 66.67% of them were located in the central domain of *16S rRNA*. Studying the effect of these mutations on the functions of *16S rRNA* molecules in protein synthesis and antibiotic resistance is of great importance.

## 1. Introduction

*H. pylori* is ubiquitous among humans,^(1)^ and one of the best studied examples of an intimate association between bacteria and humans. ^(2)^ It has infected human around 100,000 years ago (range 88,000-116,000). ^(1, 3)^ and has co-evolved with human ancestors for approximately 58000 ± 3500 years, during their first migrations from east Africa. ^(1, 4, 5)^ And it largely escaped notice until it was cultured by Marshall and Warren. ^(6)^ In fact, *H. pylori* possesses several properties that helping this bacterium to persist for several decades, transmit from generation to generation and make an intimate association with its human host. ^(2, 7)^ *H. pylori* infection is predominantly transmitted vertically from parent to child and between individuals in close contact such as in a family. ^(2, 8)^ This close transmission pattern has resulted in a clear phylogeographic differentiation within these bacteria because of the local dispersion of single nucleotide polymorphisms by high rates of homologous recombination. ^(9, 10)^ However, under poverty and inappropriate conditions, especially in the developing countries, *H. pylori* can be transmitted horizontally which makes infection with multiple strains probably more common than in the developed world. ^(11, 12)^

In Sudan, the population is culturally, linguistically and ethnically diverse with more than 597 tribes who speak more than 400 dialects and languages. ^(13, 14)^ The majority of the Sudanese population is rural with an urban population of just 33.2%, most of them are in Khartoum. ^(14, 15)^ Sudan has severely suffered war, famine and flood in recent decades and has a large population of IDPs.^(16, 17)^ Although Sudan is rich in terms of natural and human resources, the effects of the civil conflict on health, nutrition, population, the economic and social development have undoubtedly been significant. These several diverse socio-demographic factors lead to continually increasing in the prevalence rate of *H. pylori* infection ranging from 48% to 65.8% which represents a major public health challenge. ^(18, 19)^

However, studying the genetics of the *H. pylori* population has been of great interest due to its clinical and phylogeographic significance. ^(4)^ A number of studies have discovered the importance of evolutionary history to the clinical outcome of *H. pylori* infection; and indicated that the scramble of the relationship between bacterial and human ancestries at the individual level due to a consequence of migrations, invasions or racial admixture may moderate adverse outcomes for the host and disrupting the selection for a reduced virulence; and this may give some explanation to the continental enigmas of *H. pylori*. ^(12,20-23)^ Also, Phylogeny and Phylogeography of *H. pylori* strains are known to mirror human migration patterns ^(24-27)^ and reflect significant demographic events in human prehistory. ^(25, 28)^ Therefore, the phylogeographic pattern of *H. pylori* is a discriminatory tool to investigate human evolution and migration in addition to the traditional human genetic tools, e.g. mitochondrial DNA (mtDNA) and languages. ^(1)^

*16S rRNA* gene amplification and sequencing have been extensively used for bacterial phylogeny and taxonomy; and eventually, the establishment of large public-domain databases. ^(29-32)^ Several properties of the *16S rRNA* gene make it the “Ultimate molecular chronometer”,^(33)^ and the most common housekeeping genetic marker; and hence a useful target for clinical identification and phylogeny. ^(34, 35)^ These properties include: First, its present in all bacteria, often existing as a multigene family or operons, thus it is a universal target for bacterial identification.^(32, 35, 36)^ Second, the function of *16S rRNA* has not changed over a long period, so random sequence changes are more likely to reflect the microbial evolutionary change (phylogeny) than selected changes which may alter the molecule’s function. ^(33)^ Finally, the *16S rRNA* gene is large enough, approximately 1,500bp, for informatics purposes.^(34, 35, 37)^ Most importantly that the *16S rRNA* gene consists of approximately 50 functional domains and any introduction of selected changes in one domain does not greatly affect sequences in other domains i.e. less impact selected changes have on phylogenetic relationships. ^(35)^

Here in this study, two sets of primers (universal and specific for *H. pylori*) were used to amplify the *16S ribosomal* gene directly from gastric endoscopic biopsy samples collected from dyspeptic patients who had been referred for endoscopy. Sanger sequencing was performed and by matching these sequences with those available in the National Center for Biotechnology Information (NCBI) nucleotide database, we analyzed the evolutionary aspects of Sudanese *H. pylori* strains using a phylogenetic approach. The novelty of our study resides in being the first study to characterize *H. pylori* Sudanese strains detected from different regions and ethnic of Sudan using *the 16S rRNA* gene.

## 2. Material and methods

### 2.1 Study design and Study sittings

A prospective cross-sectional hospital-based study was conducted in public and private hospitals in Khartoum state from June 2018 to September 2019. These hospitals receive patients from all over the regions of Sudan. Currently, there are 16 states in Sudan which divided between four geographic regions: Eastern, Western, Northern and Central Sudan which includes the largest metropolitan area, Khartoum (that includes Khartoum, Khartoum North and Omdurman),^(38)^ see a map in Figure 3(a) for more illustration. The capital Khartoum is quickly growing and ranges between 6 and 7 million of populations, which includes approximately 2 million internally displaced persons (IDPs) from the southern war zone and the drought-affected areas in the west and east. ^(14, 17)^

**Figure 1.**
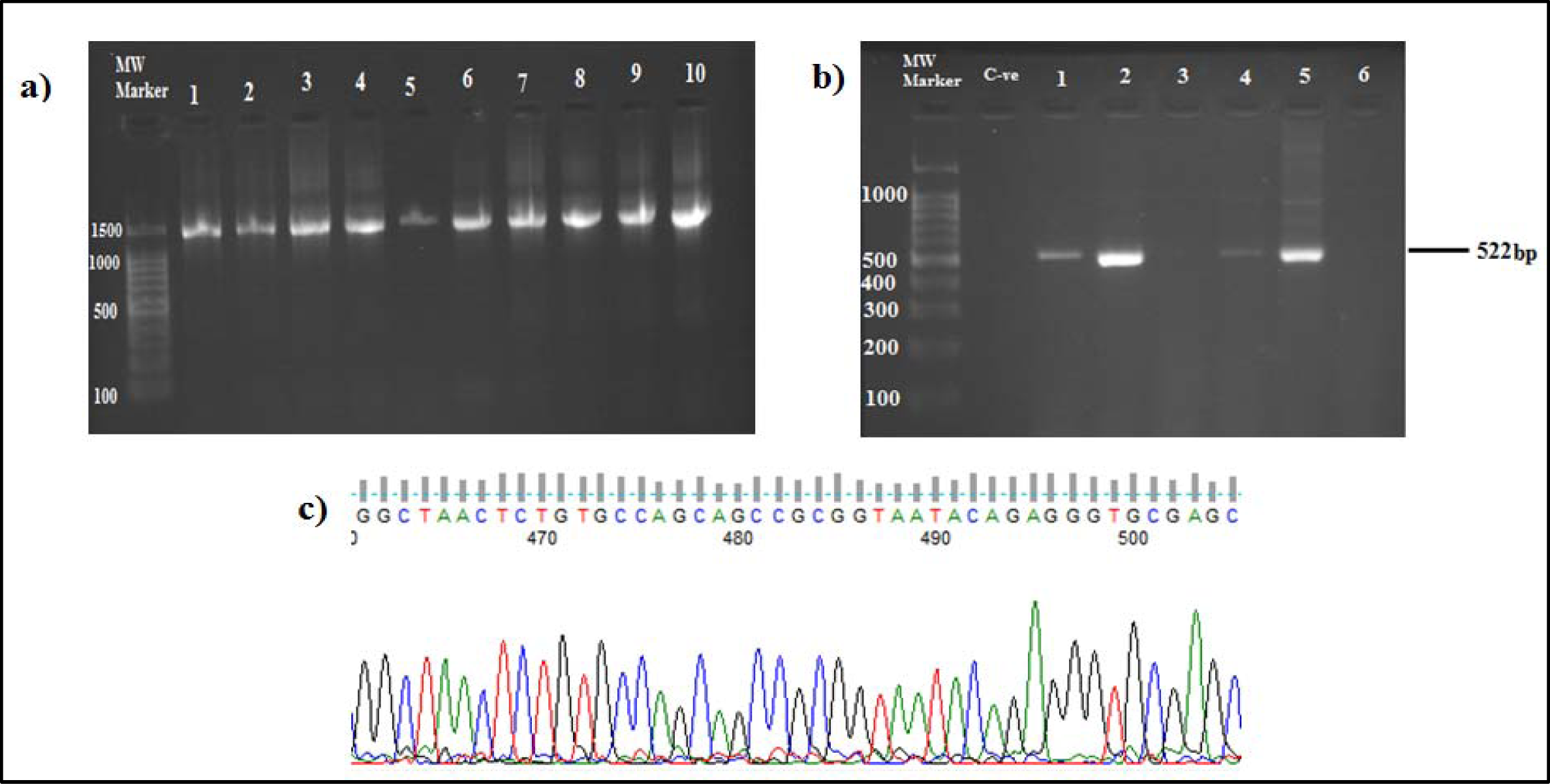
1a) and b). PCR amplification results of universal and specific H. pylori 16S rRNA gene, respectively examined on 2% agarose gel electrophoresis. 1c). Sequencing result of Acinetobacter radioresistens chromatogram using Finch TV software. The nucleotide sequence was deposited in the GenBank database under the accession number: MN845952

**Figure 2.**
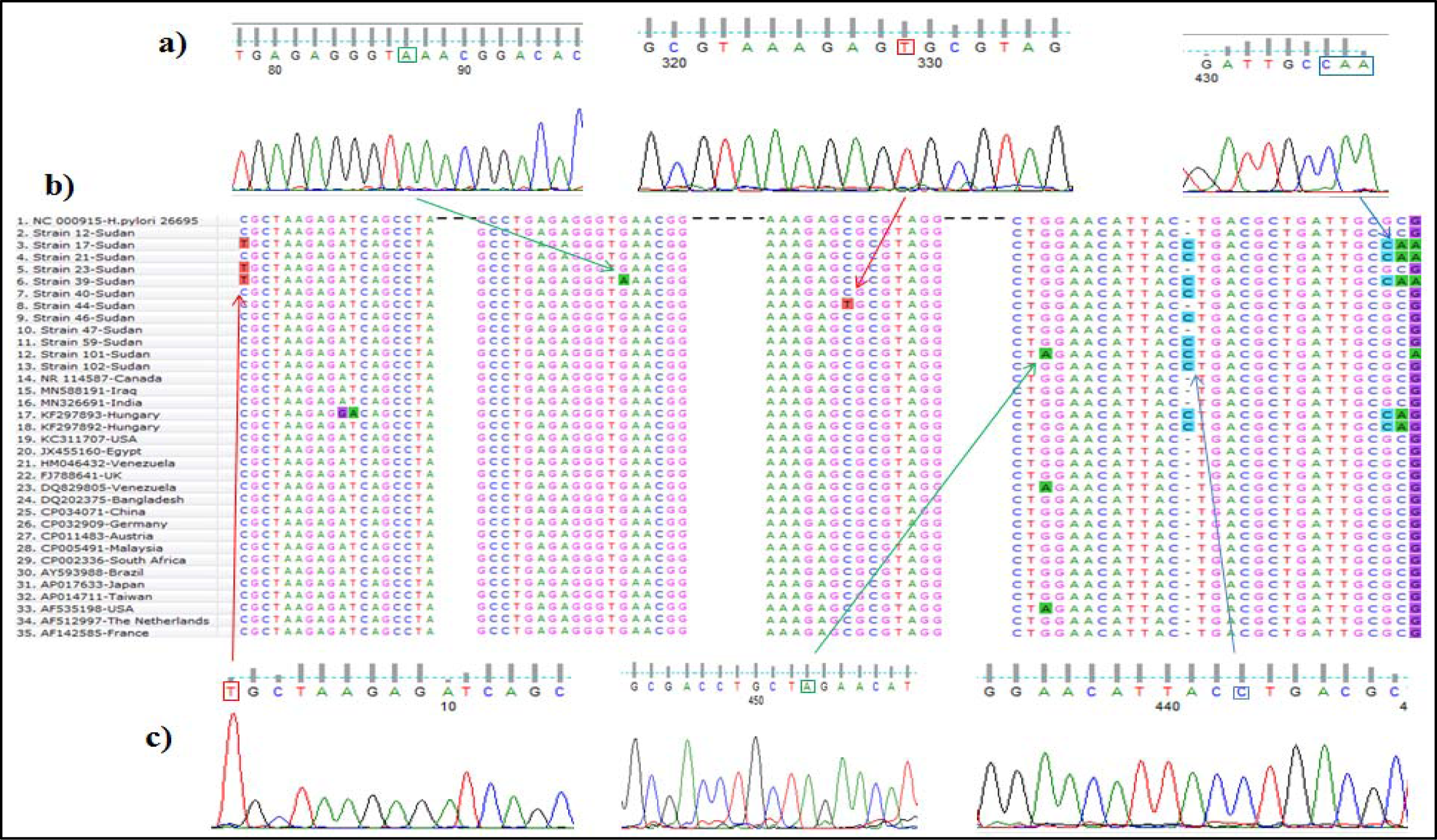
2a) and 2c) Sequencing results of chromatograms using Finch TV software show nucleotide variations in the 16S rRNA gene of H. pylori which illustrated by squares. 2b) Multiple Sequence Alignment (MSA) of 16S rRNA sequences of 12 Sudanese H. pylori strains compared with the rrnA gene of H. pylori strain 26695 (NC_000915) and other selected strains obtained from GenBank databases using Clustal W2.

**Figure 3.**
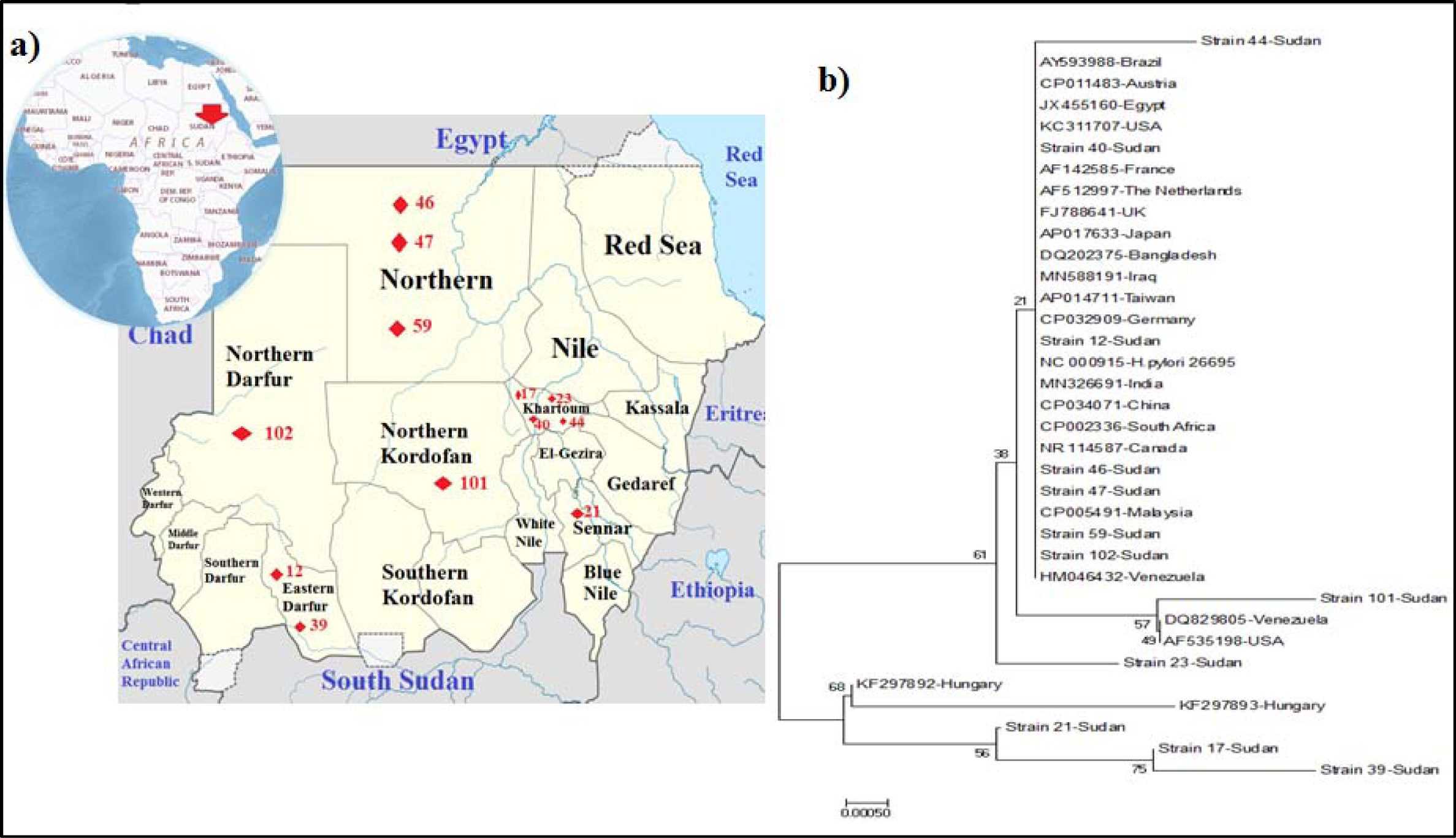
Distribution and evolutionary relationships of Sudanese H. pylori strains. 3a) Map of regional origin of strains in Sudan. 3b) The Neighbor-Joining Phylogenetic tree. The percentage of replicate trees (1000 replicates) are shown next to the branches. The evolutionary distance was computed using (JC) method and are in the units of the number of base substitution per site. Evolutionary analyses were conducted using MEGA7.

The hospitals include Ibin Sina specialized hospital, Soba teaching hospital, Modern Medical Centre and Al Faisal Specialized Hospital. All laboratory processes were performed in the Molecular biological lab at the Faculty of Medical Laboratory Sciences at the University of Khartoum.

### 2.2 Study population

The study population composed of 75 patients who had been referred for endoscopy and most of them because of dyspepsia. Structured questionnaires were provided for participants to obtain information about their socio-demographic characteristics. Patients were selected from those who were not taking antibiotics or NSAIDS. The patients gave written informed consent before they enrolled in the study. The diagnosis of gastroduodenal diseases was based on the investigation of an experienced gastroenterologist during the upper GI endoscopy procedure. While gastric cancer was diagnosed based on, in addition to gastroscopy, histology.

### 2.3 Sample collection

For DNA extraction purposes, gastric biopsies were collected in 1.5µl Eppendorf tubes with 400µl phosphate buffer saline (PBS), while for histological examination, the biopsies were transported in formalin. Then the biopsies were labeled and transported immediately to the laboratory for further processes.

### 2.4 DNA extraction

The DNA extraction was done by using the manual guanidine chloride method, as described by Alsadig *et al*. ^(39)^ Briefly, biopsies were washed with 400µl PBS and centrifuged for 5min at 100 rpm after each wash. Then, the samples were subjected to digestion by adding 400µl of WBCs lysis buffer, 200µl of 6M Guanidine chloride, 50µl of 7.5M Ammonium Acetate and 5µl of 20mg/µl proteinase K; and incubated at 37°C overnight. After that, On the following day samples were cooled down to room temperature then added 400µl of cooled pre-chilled Chloroform and centrifuged at 1000rpm for 5min. Then three layers were separated and the supernatant was collected to a new labeled Eppendorf. After that, 1ml of cooled pre-chilled absolute Ethanol was added and mixed gently back and forth quickly. Then samples were put in -20°C freezer overnight. After that, samples were subjected to quick vortex for one minute then centrifuged for 5min at 1000rpm; and the supernatants were discarded. Then washing with 70% ethanol was performed and the supernatant was drained with much care to avoid losing the DNA pellet at the bottom of the Eppendorf. Then the Eppendorf was inverted upside down of a tissue paper leaving the pellet to dry from alcohol for at least 2hours. Finally, the DNA pellet was re-suspended in 35µl of deionized water and was put into -20°C until use.

### 2.5 *16S rRNA* gene amplification

Extracted DNA was amplified for the universal *16S rRNA* gene using the following primers: (primers: F:5’-AGAGTTTGATCCTGGCTCAG-3’) (R:5’-CTACGGCTACCTTGTTACGA-3’). ^(40)^ PCR amplification was carried out with Maxime PCR PreMix Kit (i-Taq) (iNtRON BIOTECHNOLOGY, Seongnam, Korea) and a PCR thermocycler (SensoQuest, Germany). The PCR reaction mixtures contained 2.5U of i-Taq TM DNA polymerase (5U/µl), 2.5mM of each deoxynucleoside tri-phosphates (dNTPs), 1X of PCR reaction buffer (10X),1X of gel loading buffer and 1µl of DNA template. The temperature cycle for the PCR was carried out using a method described previously. ^(41, 42)^

To detect the DNA, 3µl of each PCR products was loaded onto 2% agarose gels stained with 3µl ethidium bromide (10mg/ml) and subjected to electrophoresis in 1x Tris EDTA Buffer (TEB buffer) (89mM of Tris base, 89mM Boric acid and 2mM EDTA dissolved in 1Litter H_2_O) for 30 min at 120V and 50mA. The gel was visualized under UV light illumination. A 100MW DNA ladder (iNtRON BIOTECHNOLOGY, Seongnam, Korea) was used in each gel as a molecular size standard. The amplified product for the *rrs* gene is 1500bp.

The DNA amplification for the specific *16S rRNA* gene was performed to confirm the infection of *H. pylori*. (primers: F:5’-GCGCAATCAGCGTCAGGTAATG-3’) (R:5’-GCTAAGAGAGCAGCCTATGTCC-3’). A PCR reaction was carried out using a previously described method by Abeer *et al*. ^(43)^

### 2.6 Sequencing of *H. pylori 16S rRNA* gene

The amplified *16S rRNA* gene (for the universal and specific) was purified and sequenced, using Sanger dideoxy sequencing method, commercially by Macrogen Inc, Korea.

### 2.7 Bioinformatics analysis

#### 2.7.1 Sequence analysis

The nucleotide sequence was visualized and analyzed by using the Finch TV program version 1.4.0. ^(44)^ The nucleotide Basic Local Alignment Search Tool (BLASTn; https://blast.ncbi.nlm.nih.gov/) was used for searching about the similarity with other sequences deposited in GenBank.^(45)^ The *16S rRNA* sequence was submitted in the GenBank nucleotide database under the following accession numbers: from MN845181 to MN845190 and from MN845952 to MN845954.

#### 2.7.2 Molecular Phylogenetic Analysis

Highly similar sequences were retrieved from NCBI GenBank and subjected to multiple sequence alignment (MSA) using Clustal W2 ^(46)^-BioEdit software. ^(47)^ Gblocks was used to eliminate poorly aligned positions and divergent regions of aligned sequences so the alignment becomes more suitable for phylogenetic analysis. ^(48, 49)^ The Neighbor-Joining phylogenetic tree ^(50)^ of our *16S rRNA* sequences with those obtained from the database was constructed using Jukes-Cantor (JC) model^(51)^ from Substitution (ML) model. ^(52)^ The tree was replicated 1000 replicates in which the association with taxa clustered together in the bootstrap test. ^(53)^ Molecular Evolutionary Genetics Analysis Version 7.0 (MEGA7) was used to conduct evolutionary analyses. ^(54)^

### 2.8 Statistical nalysis

Data were analyzed using GraphPad Prism 5. Regarding the prevalence of *H. pylori* infection, differences in frequency distribution by age were examined by the Mann-Whitney test. While bivariate analysis with a categorical variable was assessed by the *χ*^*2*^ test or *Fisher’s* test. The statistical significance level was determined at P<0.05.

## 3. Result

### 3.1 Characteristics of the study population

A total of seventy-five patients were included. Forty-one patients (54.67%) were male and thirty-four (45.33%) female. Forty-two patients (56%) were urban, and thirty-three (44%) were rural. The patients’ age ranged from 15 to 85 years, with a mean age of 45.11±17.45 years. Most of the participants came from northern and central Sudan. And their ethnicities were distributed as follows: Shagia (10, 13.33%), Jalyeen (9, 12%), Mahas (8,10.67%), Rezaigat (4, 5.33%), Zaghawa (4, 5.33%), Kawahla (4, 5.33%), Masalamyia (3, 4%) and Other (33, 44%). Regarding clinical symptoms of gastrointestinal disturbances of the participants, abdominal pain was the major symptom (22, 29.33%), followed by nausea (16, 21.33%). Molecular detection of *H. pylori* has shown that 28 (37.33%) of patients were positive for *H. pylori*. (Figure1(b)) Patients from western Sudan were more prone to *H. pylori* infection 50% (6/12). Bivariate analysis has found no significant differences exhibited across sociodemographic, endoscopy series and *H. pylori* infection, as illustrated in Table1 and Table2.

**Table1.** Socio-demographic characteristic of patients.

| Variables |  | Total<br>n=75 | *H. pylori* (+ve)<br>n=28 | *H. pylori* (-ve)<br>n=47 | P-value |
| --- | --- | --- | --- | --- | --- |
| Mean age (years) ± Std. Deviation (range) |  | 45.11±17.45<br>(15-85) | 41.39±17.57<br>(15-85) | 47.32±17.18<br>(21-80) | 0.1170 |
| Gender | Male | 41 (54.67%) | 15 (53.57%) | 26 (55.32%) | 1.0000 |
|  | Female | 34 (45.33%) | 13 (46.43%) | 21 (44.68%) |  |
| Residence | Urban | 42 (56%) | 15 (53.57%) | 27 (57.45%) | 0.8122 |
|  | Rural | 33 (44%) | 13 (46.43%) | 20 (42.55%) |  |
| Hospital | Public | 51 (68%) | 20 (71.43%) | 31 (65.96%) | 0.7986 |
|  | Private | 24 (32%) | 8 (28.57%) | 16 (34.04%) |  |
| History of previous infection | Yes | 42 (56%) | 15 (53.57%) | 27 (57.45%) | 0.8122 |
|  | No | 33 (44%) | 13 (46.43%) | 20 (42.55%) |  |
| The Frequency of recurrence (If Yes n=42) | One time | 32 (76.19%) | 10 (66.67%) | 22 (81.49%) | 0.4508 |
|  | More than one time | 10 (23.81%) | 5 (33.3%) | 5 (18.52%) |  |
| Family history of infection | Yes | 36 (48%) | 15 (53.58%) | 21 (44.68%) | 0.4833 |
|  | No | 39 (52%) | 13 (46.43%) | 26 (55.32%) |  |
| Smoking | Yes | 16 (21.33%) | 5 (17.86%) | 11 (23.4%) | 0.7718 |
|  | No | 59 (78.67%) | 23 (82.14%) | 36 (76.6%) |  |
| Tribe | Shagia | 10 (13.33%) | 4 (14.29%) | 6 (12.77%) | 0.5733 |
|  | Jalyeen | 9 (12%) | 4 (14.29%) | 5 (10.64%) |  |
|  | Mahas | 8 (10.67%) | 3 (10.71%) | 5 (10.64%) |  |
|  | Rezaigat | 4 (5.33%) | 1 (3.57%) | 3 (6.38%) |  |
|  | Zaghawa | 4 (5.33%) | 3 (10.71%) | 1 (2.13%) |  |
|  | Kawahla | 4 (5.33%) | 2 (7.14%) | 2 (4.26%) |  |
|  | Masalamyia | 3 (4%) | 2 (7.14%) | 1 (2.13%) |  |
|  | Other | 33 (44%) | 9 (32.14%) | 24 (51.06%) |  |
| Region | Northern Sudan | 15 (20%) | 6 (21.43%) | 9 (19.15%) | 0.2697 |
|  | Central Sudan | 31 (41.33) | 12 (42.86%) | 19 (40.43%) |  |
|  | Eastern Sudan | 3 (4%) | 2 (7.14%) | 1 (2.13%) |  |
|  | Western Sudan | 12 (16%) | 6 (21.43%) | 6 (12.77%) |  |
|  | Southern Sudan | 14 (18.67%) | 2 (7.14%) | 12 (25.53%) |  |

**Table 2.** Distribution of participants according to endoscopy series and status of *H. pylori* infection.

| Endoscopy results | No.(%) | *H. pylori*<br>(+ve) | *H. pylori*<br>(-ve) | P-value |
| --- | --- | --- | --- | --- |
| Normal gastric findings | 9 (12%) | 4 (14.29%) | 5 (10.64%) | 0.5259 |
| Esophagitis | 4 (5.33%) | 3 (10.71%) | 1 (2.13%) |  |
| Esophageal varices | 6 (8%) | 1 (3.57%) | 5 (10.64%) |  |
| Gastritis | 26 (34.67%) | 8 (28.57%) | 18 (38.3%) |  |
| Duodenitis | 6 (8%) | 3 (10.71%) | 3 (6.38%) |  |
| Peptic ulcer | 6 (8%) | 3 (10.71%) | 3 (6.38%) |  |
| Cancer | 18 (24%) | 6 (21.43%) | 12 (25.53%) |  |
| **Total** | 75 (100%) | 28 (37.33%) | 47 (62.67%) |  |

### 3.2 Analysis of *16S rRNA* sequences

12 sequences of *16S rRNA of H. pylori* from Sudanese patients were analyzed for mutations and their conservative nature, and findings revealed diversity with few differences. The description of Sudanese *H. pylori* strains is shown in Table 3. The amplicon of universal *16S rRNA* for patients 22 resulted in *Acinetobacter radioresistens* (Figure 1(c)). The patient 22 was male with 33 years and he is Mahasi from northern Sudan, and he was diagnosed with antral gastritis.

**Table 3.** Description of Sudanese *H. pylori* strains.

| Strain | Age of patient | Gender of patient | Clinical origin | Ethnic origin | Province | Region | Accession No. |
| --- | --- | --- | --- | --- | --- | --- | --- |
| Strain 12 | 28 | Female | Antral gastritis | Rezaigat | Eastern Darfur | West of Sudan | MN845953 |
| Strain 17 | 23 | Female | Distal esophagitis and antral gastritis | Halawyeen | Khartoum | Middle of Sudan | MN845187 |
| Strain 21 | 32 | Male | Antral gastritis | Kawahla | Sinnar | Southeast of Sudan | MN845181 |
| Strain 23 | 15 | Female | GRED | Manaseer | Kartoum | Middle of Sudan | MN845188 |
| Strain 39 | 58 | Male | Antral gastritis | Masalamyia | Eastern Darfur | West of Sudan | MN845189 |
| Strain 40 | 68 | Male | Duodenitis | - | Khartoum | Middle of Sudan | MN845182 |
| Strain 44 | 35 | Female | Normal gastric finding | Jalyeen | Khartoum | Middle of Sudan | MN845183 |
| Strain 46 | 52 | Male | Duodenitis | Shagia | Northern State | North of Sudan | MN845184 |
| Strain 47 | 35 | Male | GRED | Shagia | Northern State | North of Sudan | MN845954 |
| Strain 59 | 36 | Male | Normal gastric finding | Shagia | Northern State | North of Sudan | MN845190 |
| Strain 101 | 60 | Female | Gastric cancer | Zaghawa | North Kordofan | South of Sudan | MN845185 |
| Strain 102 | 56 | Male | Gastric cancer | Zaghawa | Northern Darfur | East of Sudan | MN845186 |

Regarding mutations, nucleotide variations were found at five nucleotide positions (positions 219, 305, 578, 741 and 763-764) and one insertion mutation (750_InsC_751) was present in sixty-seven percent (7/12) of our strains, the numbering was according to the *E*.*coli* sequence). ^(55)^ Three strains (17, 21 and 39) revealed a novel transition C→T at position 219. A triple base pair substitution (GCC763-764CAA) was detected in strain 17, 23 and 39. However, a cancerous strain 101 had two G → A transitions at positions 741 and 765. Strain 44, which was detected in a patient with normal gastric finding and suspected celiac disease and lives in Khartoum, had also a novel C→T transversion at nucleoside position 578. Also strain 39, which was found in a patient from west of Sudan (Eastern Darfur) and was diagnosed with antral gastritis, revealed a novel G→A transition at positions 305. See Figure 2 for more illustration.

### 3.4 Phylogenetic analysis

The phylogenetic tree diverged into two lineages. In the lineage one, all the Sudanese *H. pylori* strain clustered with strains from different countries. The *16S rRNA* sequence of strain 44, although clustered with other global strains in one clade, had a novel C→T transversion at nucleotide position 578. However, cancerous strain 101 shared a common ancestor with strains from the USA (AF535198) and Venezuela (DQ829805); and represented with them a separated clade with a bootstrap value of 57%. A novel single nucleotide variation (G→A) at position 305 of the *16S rRNA* of strain 23 made it outgroup as a kind of strain evolution with a bootstrap value of 61%.

In the second lineage, strains 17, 23 and 39 were closely related to strains from Hungary (KF297893 and KF297892). Strain 17 and Strain 39, from Khartoum and Eastern Darfur, respectively, were sisters with a bootstrap value of 75%, as presented in Figure 3.

## 4. Discussion

In this study, the phylogenetic analysis of *16S rRNA* sequences identified two lineages of *H. pylori* strains detected from different regions in Sudan which suggested differential evolution. This finding is in agreement with a number of studies that found an unusually high degree of genetic diversity in genomic sequence analyses of *H. pylori* strains in connection with their house-keeping and virulence associated genes.^(25, 56, 57)^ Strain17, Strain21 and Strain39 were derived from one lineage along with two strains from Hungary, with a bootstrap value of 61%. They shared a double base pair substitution (GC763-764CA) and one insertion mutation (751_InsC_752). However, both mutations were located in the central domain of the *16S rRNA* gene. Despite Sudanese strains were from different ethnicity and regions of Sudan but they all caused antral gastritis, while the Hungarian strains (KF297892 and KF297893) were isolated from Demodex mites and caused rosacea. ^(58)^

Laboratory diagnosis of *H. pylori* by conventional cultural methodology and phenotypic identification tests is still less specific, time consuming, increased capital costs, needs of highly skilled personnel and also it difficult to diagnose re-infections. ^(59)^ However, molecular tests and PCR technology have been raised as a useful alternative means of bacterial identification, which may circumvent some of these difficulties ^(59, 60)^ but has limited mainly by contamination and inadequate sensitivity issues. ^(61)^ In this study, molecular detection of *H. pylori* has shown that 37.33% of patients were positive for *H. pylori* in comparison to other studies conducted in different regions of Sudan, 22.2%, 59%, 65.8%, 40.1%, 48%, 21.8% by Mona *et. al*., Yousra *et. al*., Tajeldin *et. al*., Karimeldin *et. al*., Abdalsadeg *et. al*., and Mohammed *et. al*., respectively using different detection methods.^(18, 19, 62-65)^

Tetracycline is one of the 30S-targeting antibiotics that inhibit translation elongation by sterically interfering with the binding of aminoacyl-tRNA to the A-site of the ribosome.^(66)^ Therefore, mutations located in two domains (III and IV) in *16S rRNA*: helix 34 and the loop next to helix 31^(67)^ can affect the conformation of the tetracycline-binding site, leading to high-level resistance. ^(66)^ In this study, six mutations were detected in two domains (I and II) which are in regions of the *16S rRNA* not closely associated with either tetracycline or tRNA binding sites. This finding is partially in agreement with a study conducted by Catharine *et. al*., that found six nucleotide changes in two tetracycline-resistance strains. Two of these changes were located in domain I and domain II, G360A and deletion of G771, respectively. ^(68)^ However, we observed that 66.67% of nucleotide variations were located in the central domain of *16S rRNA* (nucleotides 567–915) with five associated ribosomal proteins (S15, S6 and S18) folds into the platform of the small subunit. ^(69-72)^ Moreover, deleterious mutations (C18G, A55G, A161G, A373G, G521A, C614A, A622G, and deletion of one A in a triple-A cluster 607–609) in domains I and II which were suggested by Aymen *et. al*., in purposes of revealing covered putative functional regions of the ribosome and aiding in the development of new antibiotics,^(73)^ were found conserved in our strains, as illustrated in Figure 2(b).

Interestingly, the insertion mutation (InsC), that was detected in sixty-seven percent (7/12) of our strains, between two nucleotides (750 and 751) located at the lower portion of helix 22 (H22), which is one of the lower three-helix junctions (3HJ) of *16S rRNA* that bind with S15.^(74, 75)^ Also, the mutation in cancerous strain 101 (G → A transition at positions 741) was located in the upper portion of helix 22 (H22). The molecular dynamic (MD) simulations of Wen *et. al*., showed that *16S rRNA* and S15 bind across the major groove of H22 via electrostatic interactions, i.e. the negatively charged phosphate groups of G658, U740, G741 and G742 bind to the positively charged S15 residues Lys7, Arg34 and Arg37. ^(74)^ However, studying the effect of these mutations on the functions of 16S rRNA molecules in protein synthesis and antibiotic resistance is of great importance especially for essential regions like the central domain. ^(76)^ For example, Prescott *et. al*., in 1990 found that the presence of substitution from a C to G at position 726 induces the synthesis of heat shock protein and affects the expression levels of various proteins. ^(77)^ Also, mutations in the sequence 911-915 were conferred Streptomycin-resistance by impaired the binding of this antibiotic.^(78-80)^

We acknowledge that limitations of the study are the small sample size and the phylogenetic tree was built based on the *16S rRNA* gene only which is unable to cover more complex evolutionary events and distinguish between closely related strains or species. ^(14)^ Therefore, further studies with a large sample size and building a phylogeny by increasing the number of genes analyzed like MLST.

## In conclusion

The phylogenetic analysis of *16S rRNA* sequences identified two lineages of *H. pylori* strains detected from different regions in Sudan. Sex mutations were detected in regions of the *16S rRNA* not closely associated with either tetracycline or tRNA binding sites. 66.67% of them were located in the central domain of *16S rRNA*. Studying the effect of these mutations on the functions of *16S rRNA* molecules in protein synthesis and antibiotic resistance is of great importance.

## Declarations

### Ethical approval and consent to participate

Approval to conduct this study was obtained from the Khartoum ministry of the health research department, University of Khartoum and Research Ethics Committees of hospitals. The patients gave written informed consent before they enrolled in the study.

### Consent for publication

All authors consent for publication

### Availability of data and materials

All data generated or analyzed during this study are included in this published article

### Authors’ contributions

MAH and EMI supervised the methodology and revised the manuscript. ABI, HGH, LBI, AMA and MMAI collected the samples. ABI, MASA and SME extracted the DNA. ABI, MASA and HNA amplified the *16S rRNA* gene. ABI analyzed the data and wrote the manuscript. MAH edited and revised the final manuscript.

## Acknowledgments

We would like to thanks patients who participated in the study and the staff of the gastroscopic unit in Ibin Sina specialized hospital, Soba teaching hospital, Modern Medical Centre, and Al Faisal Specialized Hospital.

